# Genetic changes of *P. vivax* tempers host tissue-specific responses in *Anopheles stephensi*

**DOI:** 10.1101/774166

**Authors:** Sanjay Tevatiya, Seena Kumari, Charu Chauhan, Deepak Singla, Tanwee Das De, Punita Sharma, Tina Thomas, Jyoti Rani, Kailash C Pandey, Veena Pande, Rajnikant Dixit

**Affiliations:** Laboratory of Host-Parasite Interaction Studies, ICMR-National Institute of Malaria Research, Dwarka, New Delhi-110077, India; Department of Biotechnology, Kumaun University, Nainital, Uttarakhand, India

## Abstract

In our preceding study (Sharma et al., 2019; *BioRxiv*) we showed that in the gut lumen *Plasmodium vivax* follows a unique strategy of immuno-suppression by disabling gut flora proliferation. Here, we further demonstrate that post gut invasion, a shrewd molecular relationship with individual tissues such as midgut, hemocyte, salivary glands, and strategic changes in the genetic makeup of *P. vivax* favors its survival in the mosquito host. A transient suppression of ‘metabolic machinery by early oocysts, and increased immunity’ against late oocysts suggested a unique mechanism of gut homeostasis restoration and *Plasmodium* population regulation. Though a hyper immune response of hemocyte was a key to remove free circulating sporozoites, but a strong suppression of salivary metabolic activities, may favor successful survival of invaded sporozoites. Finally, genetic alteration of *P. vivax* ensures evasion of mosquito responses. Conclusively, our system-wide RNAseq analysis provides first genetic evidences of direct mosquito-*Plasmodium* interaction and establishes a functional correlation.

**Author Summary:** Malaria transmission dynamics is heavily influenced by mosquito –parasite interaction. When passing through tissue specific barriers, *Plasmodium* have to compromise by losing its own population, but genetic relation is unknown. To win the developmental race *Plasmodium* need to overcome two important immuno-physiological barriers. First one accounts an indirect 24-30hr long pre-invasive gut-microbe-parasite interaction in the gut lumen. And second one follows a direct post gut invasive 14-18 days interaction with midgut, hemocyte and salivary glands. During pre-invasive phase of interaction, we showed *Plasmodium vivax* follows immuno-suppression strategy by restricting microbial growth in the gut lumen. Here, we demonstrate that switch of parasite from one stage to another stage within mosquito vector is accompanied by genetic changes of parasite. Our data suggests genetic makeup change enables the parasite to manipulate the metabolism of mosquito tissues. This strategy not only clear off multifaceted mosquito’s tissue specific immune responses, but also favors *Plasmodium* own survival and transmission. Comprehending this tissue specific interaction between host and parasite at molecular level could provide new tool to intervene the plasmodium life cycle within vector.

## Introduction

Evolution and adaptation of adult female mosquitoes to blood feeding have significant influence on their reproductive outcome and disease transmission dynamics. Sexual cycle of malaria causing *Plasmodium* immediately begins after ingestion of gametocytes containing blood meal by *Anopheline* mosquitoes (Bennink, Kiesow, & Pradel, 2016, Kuehn & Pradel, 2010, Talman, Domarle, McKenzie, Ariey, & Robert, 2004). Within the lumen of midgut, male and female gametocytes fuse to form zygote which then transformed into motile ookinete (Aly, Vaughan, & Kappe, 2009)(Bennink et al., 2016). Eventually, within 24hrs transformed ookinetes traverse through the gut epithelium to reach basal lamina either through intracellular and/or intercellular routes and transform to tiny oocysts (Baton & Ranford-Cartwright, 2004) (Han, Thompson, Kafatos, & Barillas-Mury, 2000). During this phase mosquito imparts early defense response through nitration of the midgut and activation of signaling pathways (Ramphul, Garver, Molina-Cruz, Canepa, & Barillas-Mury, 2015). The midgut nitration results in modification of ookinete surfaces, which makes them perceptible to mosquito complement like system (Garver, de Almeida Oliveira, & Barillas-Mury, 2013)(Shiao, Whitten, Zachary, Hoffmann, & Levashina, 2006). Whereas signaling pathways show different response towards different *Plasmodium* species, such as IMD pathway works more effectively against *P. falciparum* than *P. bergei* and Toll pathway is more responsive against *P. bergei*, and *P. gallinaceum* (Kumar, Gupta, Yeon, & Barillas-Mury, 2004)(Cirimotich, Dong, Garver, Sim, & Dimopoulos, 2010). Active pathways boost rapid production of immune peptides i.e. AMPs (Cecropin, gambicin etc.) which are responsible for parasite killing. Majority of ookinetes killing is targeted at the basal side of midgut epithelium, through an indirect hemocyte mediated cellular response such as lysis or melanization (Smith et al., 2016; Belachew, 2018). Recent study shows, during immune active complement system the hemocyte responds through micro vesicles. (Castillo, Barletta Ferreira, Trisnadi, & Barillas-Mury, 2017). Once reached to basal lamina, the surviving ookinetes rapidly transform to oocysts. These hidden young oocysts gradually mature to round shaped large oocysts and remain protected from any external response until burst up into millions of sigmoid shaped sporozoites. The underlying mechanism that how maturing oocysts (i) became resistance to any external response; (ii) regulates their time dependent developmental transformation to mature oocysts; (iii) trigger sporozoites release outside the basal lamina by rupturing the oocysts, is not well known.

Once released, millions of free circulatory sporozoites (*fc*Spz) are rapidly killed by the immune blood cells i.e. hemocytes. Despite of a substantial reduction in the population of sporozoites (∼ 85 Fold),(Gouagna et al., 1998), a significant number of parasites succeed to specifically invade the salivary glands and survive within it. Several hemocyte mediated mechanisms such as melanization, lysis, engulfment, toxic anti-*Plamsodium* immune effector molecules have been proposed to kill the free circulating sporozoites, but the molecular nature of these interactions is unknown (De et al., 2018; Julián F. Hillyer, Schmidt, & Christensen, 2003; Smith et al., 2016). Even for the salivary invasion *Plasmodium* needs salivary gland proteins such as saglin, MAEBL, PCRMP (Kariu, Yuda, Yano, & Chinzei, 2002). Recent study also shows that *Plasmodium* invasion induced several salivary immune proteins such as SP14D1, Ficolin, SP24D, PGRP, LRIM (Roth et al., 2018). However, the genetic basis that how sporozoites (i) manage to avoid hemocyte mediated killing responses; (ii) guide its movements for salivary invasion; and (iii) promote its virulence is not well understood (Mueller, Kohlhepp, Hammerschmidt, & Michel, 2010; Roth et al., 2018).

In our preceding study (see Sharma et al., 2019; *BioRxiv*) we demonstrate that in the gut lumen how *Plasmodium vivax* follows a unique strategy of immuno-suppression by disabling gut flora proliferation. Here we further decoded the post gut invasive molecular complexity of tissue specific-parasite interactions. A comprehensive *RNA*-*Seq* analysis of *P. vivax* infected midgut, hemocyte and salivary glands not only identified mosquito transcripts, but also captured a large pool of *P. vivax* transcripts expressing in respective tissues. A detailed annotation and transcriptional profiling revealed that *P. vivax* has ability to avoid immuno-physiological responses by altering its own genetic architecture. Manipulating tissue specific immuno-physiology of the mosquitoes may render the *Plasmodium* development and hence the transmission.

## MATERIALS AND METHODS

### Mosquito Rearing, Tissue specific *RNA-Seq* library sequencing and analysis

As described in our preceding study, essentially we followed same artificial membrane feeding assay (AMFA) protocol to infect mosquitoes with *P. vivax*, RNA isolation and *RNA-Seq* library preparation and sequencing (Sharma et al., 2019; *BioRxiv*). Briefly, post confirmation of *P. vivax* infection, ∼ 20-25 mosquitoes were dissected at 3-6Days Post Infection (DPI) and 8-10 DPI for midgut, 9-12DPI for hemocytes and 12-14DPI for salivary glands in PBS and collected in trizol reagent. Total RNA was isolated from each dissected tissue sample, and respective double stranded cDNA library was prepared using well established protocol, described earlier (De et al., 2018; Dixit et al., 2011; Sharma et al., 2015). The sequencing of whole transcriptomes was performed on Illumina NextSeq. Post filtration of adaptor sequences and low quality (QV < 20) reads, high quality clean reads were used to make assembly using Trinity software. From assembled contigs, CDS were predicted from longest reading frame using Transdecoder. Subsequently, the predicted CDS were annotated using BLASTX against NCBI NR database. For heat map and Venn diagram analysis online softwares (Lam et al., 2016) were used. Technical plan and complete workflow strategy is shown in Fig. 1 (Panel A).

**Fig1.**
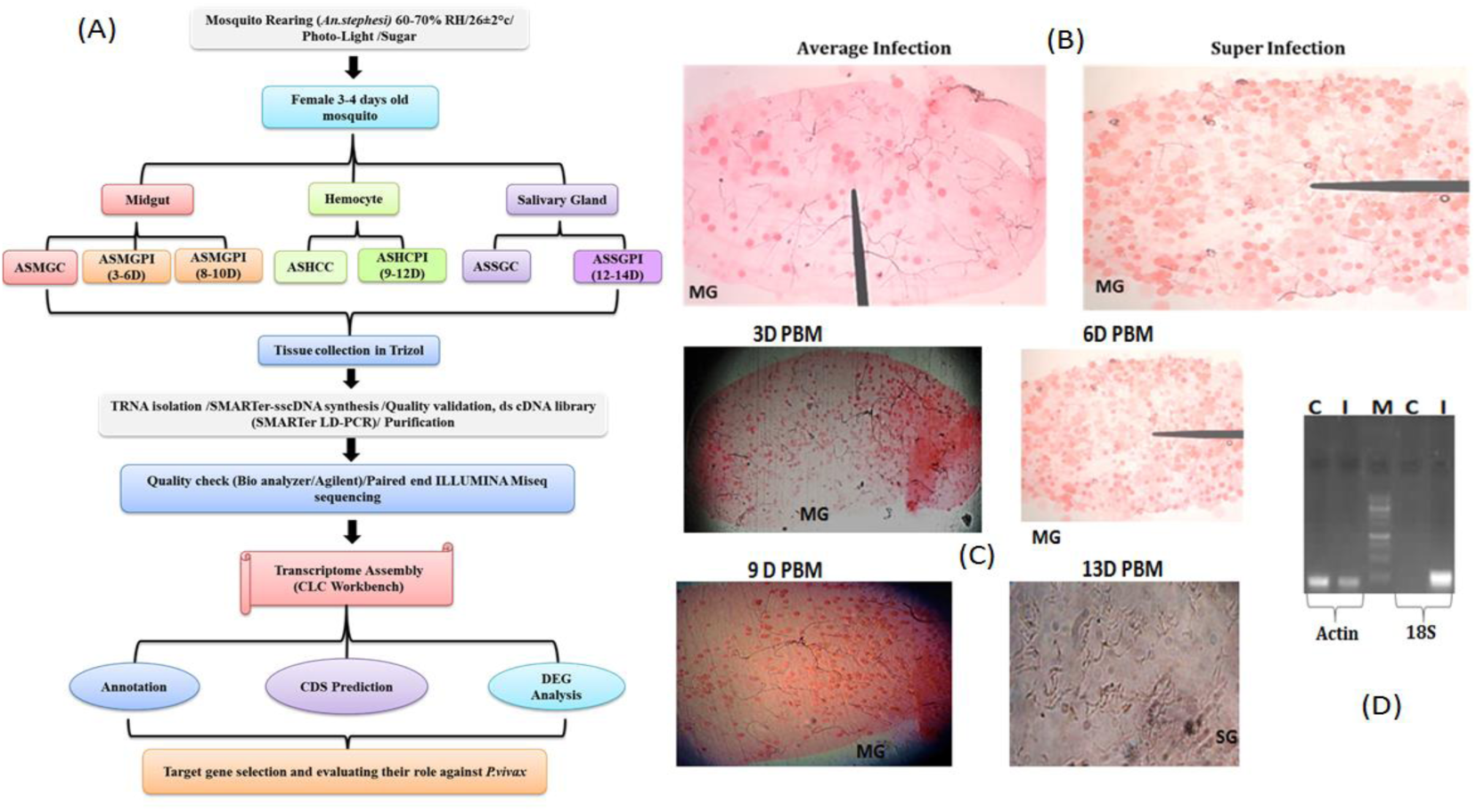
Technical design and mosquito tissue collection strategy for tissue specific RNAseq analysis: <u>**Panel (A)**</u>: Schematic presentation of designed technical work plan; <u>**Panel (B)**</u>: a comparative microscopic analysis of average infected and super infected mosquitoes midgut, six days’ post *Plasmodium vivax* infection; <u>**Panel (C)**</u>: Microscopic view of successful parasite development in the mosquito *Anopheles stephensi* where infected tissues include: midgut (MG) & Salivary Glands (SG) were collected in PBS. Post blood meal (PBM), the confirmed infected mosquitoes MG were dissected at different time interval e.g. 3-6 days, where dark Red Circle indicate the oocysts in the individual midgut. Spindle shape bodies in the last image of Panel (C) indicate sporozoites observed in the salivary glands collected 13 DPI; <u>**Panel (D)**</u>: Molecular confirmation of parasite infection in the mosquito by RT-PCR and agarose gel electrophoresis: Uninfected (C) & Infected (I) midgut cDNA samples were subjected to 28 PCR cycle amplification using mosquito specific Actin & parasite specific 18S primers. M=Molecular marker. Note: second image in panel B i.e. Super infection is same in panel C representing 6D PBM oocysts infection

### GO annotation, molecular cataloging and gene expression profiling

A comprehensive GO annotation, molecular cataloging and functional prediction analysis was performed using BLAST2GO software (Conesa, Götz, García-gómez, Terol, & Talón, 2005). For immune cataloging all predicted CDS were subjected for BLASTX analysis against *An. gambiae, D. melanogaster, Ae. aegypti* and *Cu. pipiens* Immunome database (http://cegg.unige.ch/Insecta/immunodb), as described earlier (Thomas et al., 2016). Then transcripts having E-value <e^-10^ were shortlisted, catalogued and compared to select transcripts for expression profiling. For cataloging tissue specific *P. vivax* genes, from pre-analyzed whole transcripts database (NR/BLASTX), the best match *Plasmodium* transcripts were retrieved as FASTA file. A standard GO annotation was performed and *P. vivax* transcripts were analyzed for functional prediction. The gene expression analysis of the selected mosquito and/or *P. vivax* transcripts was monitored as described earlier (De et al., 2018; Thomas et al., 2016); also see Sharma et al., 2019; *BioRxiv*)

## Results

### Working hypothesis development and RNAseq data generation

Previous studies targeting individual tissues either midgut or salivary glands, have been valuable to understand the mosquito-parasite interactions. But still, there are several unresolved questions that how (i) each tissue viz. midgut, hemocytes and salivary glands together coordinate and manage the challenge of *Plasmodium* infection; and (ii) how *Plasmodium* avoid the tissue specific responses for its survival and transmission. To partly answer and resolve tissue specific molecular complexity, we developed a working hypothesis. We opined when blood meal itself significantly alters mosquitoes ‘metabolic physiology, *Plasmodium* infection may cause an additional burden of immune activation. Thus, mosquitoes may need to follow a dual management strategy. Alternatively, for its survival parasites may also suppress/misguide the host tissues responses. (See Sharma et al., 2019; *BioRxiv*/Fig.1).

To test and evaluate the hypothesis we designed a strategy to capture molecular snapshot of the three tissues directly interacting with *Plasmodium vivax.* After establishment of artificial membrane feeding protocol, we fed mosquitoes with clinically diagnosed *P. vivax* infected patient’s blood sample (0.5-2% gametocytemia). In our regular experience, we noticed average infection intensity of 50-130 oocysts/midgut. But in a few experimental studies, we noticed a super infection of *P. vivax*, raising average infection intensity to 300-380 oocysts/midgut (Fig.1; see Panel B). Thus, for our transcriptomic study we specifically targeted super infected mosquito group with a possibility of retrieving *P. vivax* transcripts.

From infected 20-25 individual mosquitoes, we dissected and pooled targeted midgut, hemocytes, and salivary glands tissues and performed a tissue specific comparative *RNAseq* analysis. As per technical design, we sequenced two midgut samples covering early (3-6 days) pre-mature to maturing oocysts; and late (8-10 days) fully matured/bursting oocysts; one hemocyte sample pooled from 9-12 days covering free circulatory sporozoites; and one sample of salivary glands 12-14 DPI (Fig.1; see Panel A). For comparative study, we sequenced the naïve blood fed mosquitoes’ tissues collected from the same aged mosquitoes. In case of midgut we sequenced only one sample 3-4 days post blood meal (also see Sharma et al., 2019; *BioRxiv*). From the seven *RNAseq* libraries we generated and analyzed total of 28.5 million reads (ST1). Table-1 represents complete stat of the sequencing and analysis for each tissue specific *RNAseq* data.

**Table-1:** Complete sequencing stat of tissue specific RNAseq libraries.

| Sl. No. | Sample Name | No. of Reads | No. of Contigs | No. of Transcripts | Total BLAST Hits (NR) |
| --- | --- | --- | --- | --- | --- |
| 1. | AS-MG-C | 4673408 | 74,548 | 11159 | 10219 (91%) |
| 2. | AS-MG-Pv. Inf (3-6D) | 3625905 | 52,187 | 12065 | 11011 (91%) |
| 3. | AS-MG-Pv. Inf (8-10D) | 5399936 | 42,212 | 7585 | 7222 (95%) |
| 4. | AS-HC-C | 3827903 | 30,959 | 6236 | 6044 (96%) |
| 5. | AS-HC-Pv. Inf | 3235689 | 18,462 | 3492 | 3333 (95%) |
| 6. | AS-SG-C | 3782999 | 64,960 | 7674 | 6840 (89%) |
| 7. | AS-SG-Pv. Inf | 4396363 | 32,110 | 4492 | 4100 (91%) |

### *Plasmodium vivax* alters the molecular architecture of mosquito tissues

To understand that, how *P. vivax* infection influences the tissue specific molecular responses; we compared global change in the transcript abundance of ‘uninfected’ and ‘infected’ blood fed mosquito tissues. Our initial attempt of mapping of cleaned reads (filtered low quality, microbial and *P. vivax* origin) to the available draft reference genome was unsuccessful. Alternatively, we mapped all the high quality reads against *denovo* assembled reference map, as described earlier (De et al., 2018; Sharma et al., 2015; Thomas et al., 2016). Though, a read density map analysis revealed *P. vivax* infection causes a significant suppression in the salivary glands and midgut transcripts, but cause a greater shift in the read density of the infected hemocyte transcripts **(Fig.2)**. To further uncover the molecular and functional nature of the encoded proteins, we carried out a comprehensive GO annotation, cataloged and extensively profiled tissues specific shortlisted genes altered in response to *P. vivax* infection.

**Figure2:**
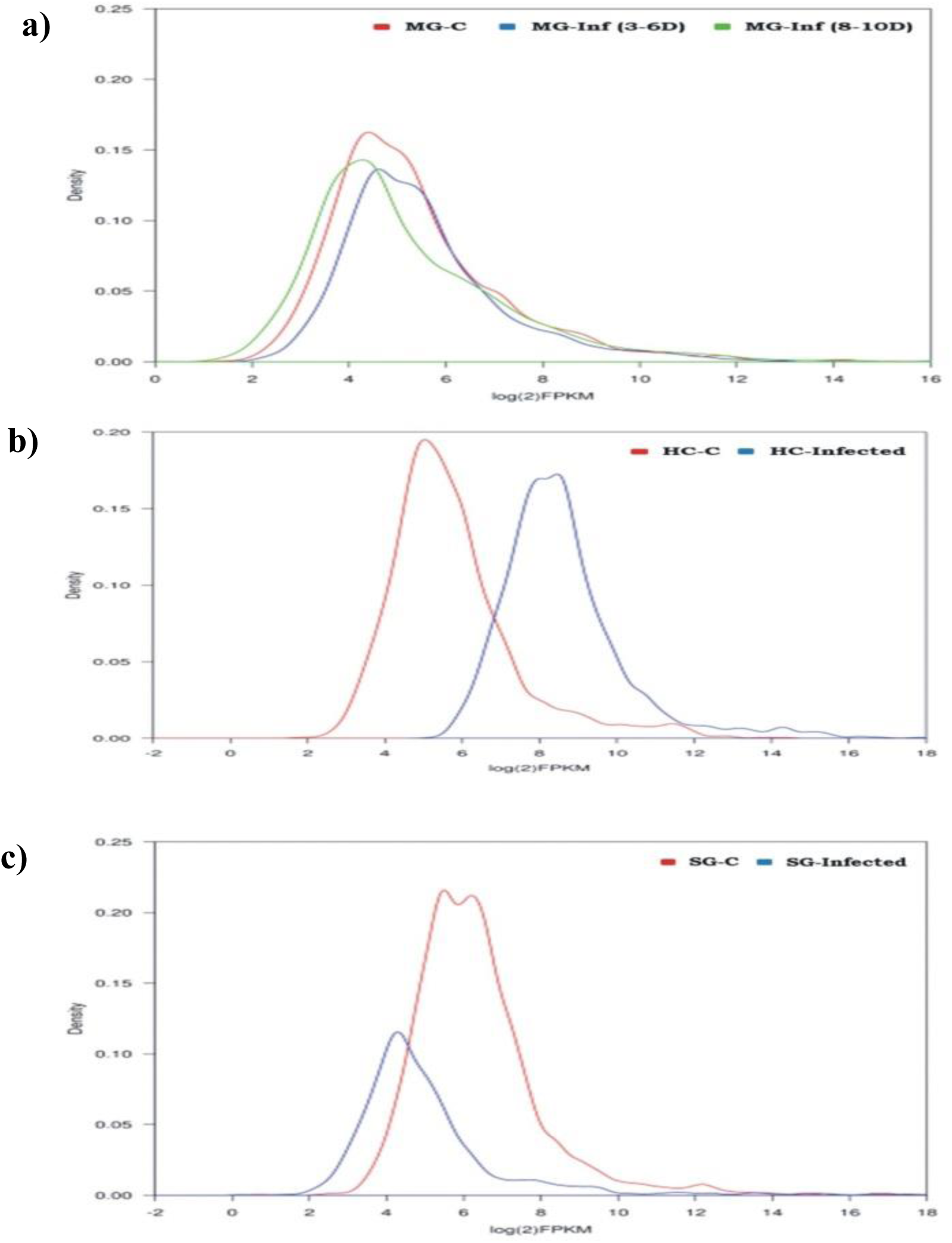
Read density maps showing change in global gene expression pattern during *P.vivax* infection: **(a)** when compared to blood fed midgut control (MG-C), the *P. vivax* infection caused slight suppression of midgut genes during early i.e. 3-6 DPI (MG-Inf 3-6D) and then regained its normal homeostasis during late i.e. 8-10 DPI (MG-Inf 8-10D);**(b)** *P.vivax* infection caused major shift in the global gene expression of hemocytes (HC-Infected) when compared with blood fed hemocytes (HC-C); **(c)** Salivary gland genes showed suppression during parasite infection (SG-Infected).

### Midgut response to *Plasmodium* oocysts development

A comparative catalogue of biological process (L4) unraveled that an early maturing oocyst development coincides with the suppression of majority of midgut proteins(ST1). Though oocyst development exceptionally induces the expression of proteins encoding cellular catabolic, organic substance catabolic and organic acid metabolic processes, but also restricts expression of proteins having regulatory functions (Fig. 3a). Interestingly, when compared to naïve midgut, late stage oocysts showed a re-enrichment of the gut proteins having common functions (Fig.3a). Venn diagram analysis of annotated transcripts showed that 877, 1036 and 1251 transcripts are uniquely restricted their expression to naïve, early and late oocysts infected mosquito guts, respectively (Fig.3b). A heat map analysis of differentially expressed common transcripts showed a significant alteration in response to early oocysts infection (Fig.3c). Taken together, these observations suggested that fast developing oocyst of *P. vivax* may cause a biphasic modulation of gut metabolic machinery, allowing early suppression and late recovery for the maintenance of homeostasis.

**Figure 3:**
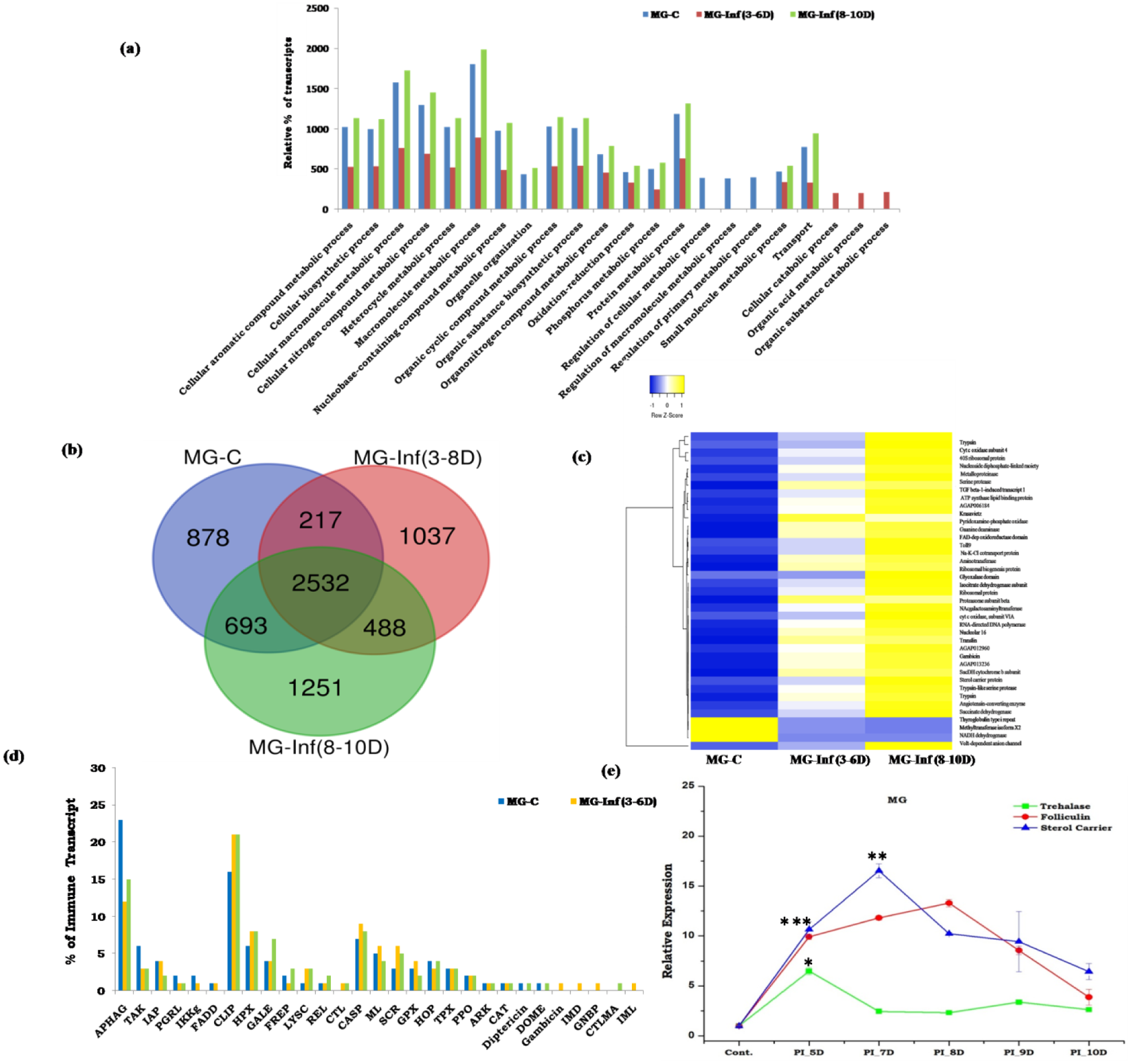
Detailed molecular cataloguing of midgut transcriptome in response to *P. vivax* infection: **(a)** Comparative Gene Ontology (GO-biological process/Level 4) analysis of midgut transcriptome revealed an initial suppression/restricted expression of most of the midgut genes during early oocyst infection (MG-Inf 3-6D) followed by re-enrichment of the respective transcripts at later stages i.e. 8-10 DPI (MG-Inf 8-10D); **(b)** Venn diagram analysis represents, genes expressing uniquely in midgut after blood meal, during early oocyst infection as well as matured oocyst infection; **(c)** Heat map showing differential expression of the common midgut transcripts in response to *P. vivax* infection; **(d)** Comparative gut immunome analysis showing alteration of immune proteins of common function as well exclusive increased percentage of immune proteins (CTL: C-type lectin; Gambicin; GNBP: Gram-negative bacteria-binding proteins; IMD: Immune deficiency pathway member; CTMLA: C-type lectin mannose binding group and IML: Immunolectins) in response to early oocyst infection; **(e)** Time dependent relative gene expression analysis of selected gut transcripts: Trehalase p<0.01; Folliculin p<0.007 and Sterol carrier p<0.002 (S2_primer list) encoding nutritional responsive proteins in response to *P. vivax* infection.

To further clarify that how *Plasmodium vivax* infection alters gut immuno-physiological responses, we identified, catalogued and compared expression of selected transcripts in response to blood meal and *P. vivax* infection. A comparative gut-immunome analysis showed an increased percentage of immune-transcripts, except a few classes such as Autophagy, TAK, IAP, PGRP, IKKG (Fig.3d). Additionally, we also observed an exclusive enrichment of several classes of immune proteins such as CTL, Gambicin, GNBP, IMD, CTMLA and IML in response to early infection (ST2). As expected, blood meal causes a transient change in immune gene expression (S1), but we observed a multifold enrichment of Gambicin expression after 48 hours and late induction of other AMPs such as C1, C2 and D1 (see Fig.8/ Sharma et al., 2019; *BioRxiv*). Together these data suggested a time dependent action of distinct AMPs against *P. vivax* is necessary to delimit the gut specific oocysts development. Furthermore, a rapid induction of transcripts encoding Folliculin, Trehalase and Sterol carrier, demonstrated their possible role to manage nutritional imbalance, during *P. vivax* development (Fig. 3e).

### Hemocyte response to free circulatory sporozoites

Infected mosquito hemocytes showed a slight enrichment of transcripts (1408) encoding diverse nature proteins(ST1). The expression of 1128 transcripts linked to organelle organization and ribonucleoprotein complex biogenesis functions remains uniquely associated to infected hemocytes (Fig. 4 a,b). Interestingly, we observed at least 966 transcripts encoding proteins of cellular catabolic and organic acid metabolic processes, remained restricted to naïve blood fed mosquito hemocytes. Surprisingly, opposite to this in the gut, similar categories of proteins were exclusively induced in response of early oocysts infection (see Fig. 3, 4). FPKM (Fragments Per Kilo base of transcript per Million mapped reads) based heat map analysis further showed a unique modulation of common transcripts, having restricted expression either in the naïve or infected mosquito hemocytes (Fig. 4c).

**Figure 4:**
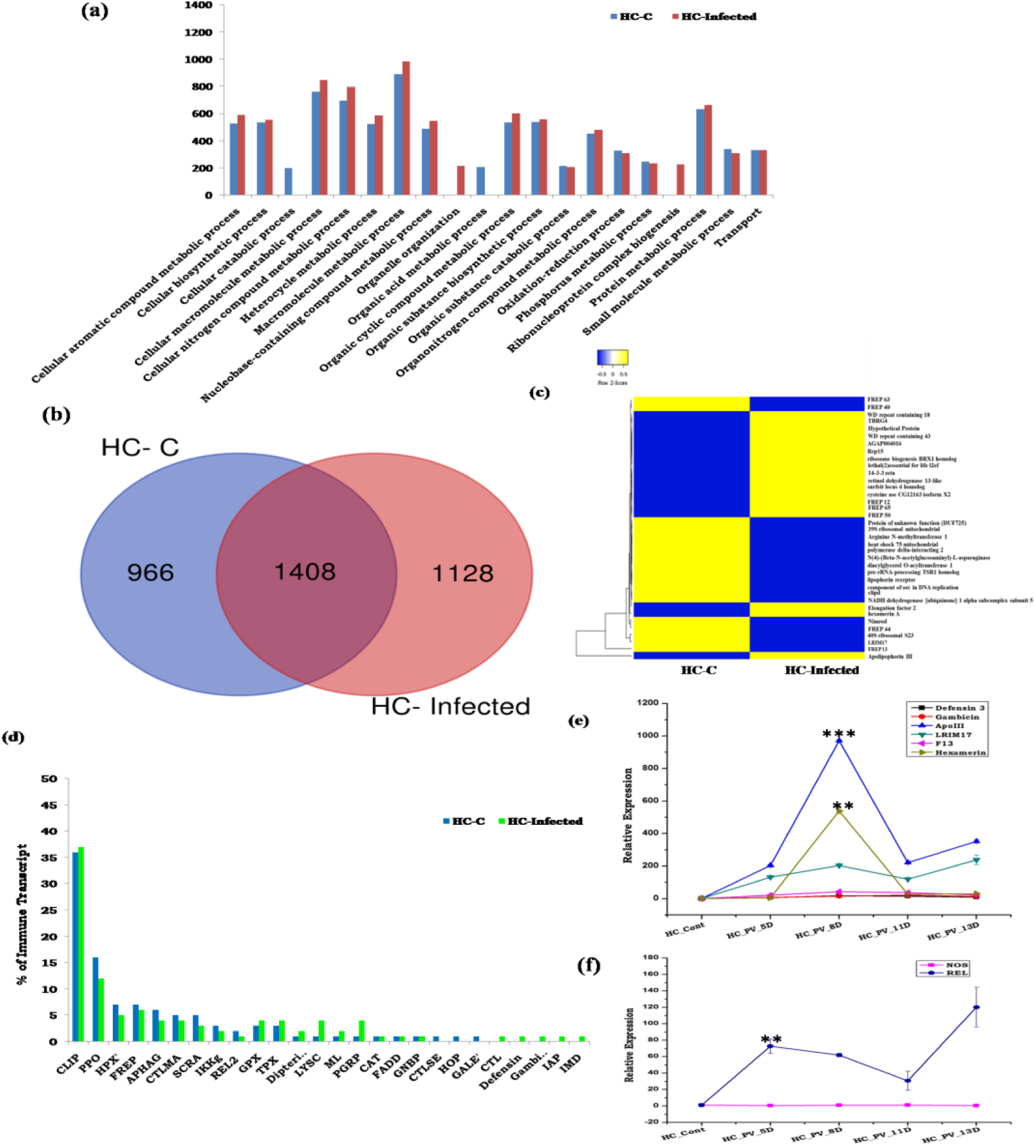
Comparative analysis of molecular architecture of hemocytes transcriptome during *P. vivax* infection: **(a)** Gene ontology (GO-biological process/Level 4) analysis of hemocyte transcriptome showing exclusive enrichment of transcripts having role in organelle organization and ribonucleoprotein complex biogenesis in response to *P. vivax* infection; **(b)** Venn diagram analysis showing 1128 transcripts expressing exclusively in *P. vivax* infected hemocytes (HC-Infected), while 966 genes expression restricted to blood fed hemocytes only (HC-C); **(c)** FPKM based heat map expression analysis of common transcripts which are either restricted to blood fed or *P. vivax* infected hemocytes; **(d)** Comparative catalogue of immune transcripts from uninfected and parasite infected hemocytes revealing exclusive enrichment of unique protein families (CTL: C-type lectin; Gambicin; Defensin; IMD: Immune deficiency pathway member; and IAP: Inhibitors of apoptosis) in response to *P. vivax* infection**;(e)** Transcriptional response of selected AMPs (Antimicrobial Peptides) and Non AMPs (Apo III p<2.06E-06, hexamerin p<.0002 and FREP13: Fibrinogen related Protein 13) in the hemocytes after 5, 8, 11 and 13 DPI with *P. vivax*; **(f)** Transcriptional profiling of signaling molecules (REL: Relish p<.003; and NOS: Nitric oxide synthase) in *P. vivax* infected hemocyte samples (S2_primer list).

Since mosquito hemocytes mount a highly specific cellular immune response, next we targeted to decode the molecular nature of immune interaction between mosquito hemocytes and free circulating sprorozoites (*fc*SPZ) of *P. vivax*. A comparative immunome(ST2) analysis indicated that *fc*SPZ causes a greater suppression of many immune family proteins, except the marked enrichment of GPX, TPX, Dipetricin, LYSC, PGRP and ML family proteins (Fig. 4d). Interestingly, like midgut, infected mosquito hemocytes also showed an exclusive induction of similar classes of immune proteins such as CTL, Gambicin, Defensin, IAP and IMD. (Fig.4d). A significant enrichment of REL than NOS coincided with early oocysts development in gut (Fig.4f). We also observed an exclusive induction of physiologically active non-AMPs class of immune proteins such as ApoIII, hexamerin and FREP13 during 8^th^ day of *P. vivax* infection.

### Salivary glands response to *P. vivax* infection

To the contrast of midgut and hemocytes, *P. vivax* infection impaired the molecular architecture of the salivary glands by delimiting the expression of common salivary transcripts(ST1). Additionally, we also observed a restricted expression of primary metabolic process proteins in uninfected mosquito salivary glands (Fig. 5a). A venn diagram analysis further indicated that only 1427 (28%) annotated salivary proteins shared a common function. Whereas, a large pool of 3069 (60%) transcripts are restricted to uninfected mosquito salivary glands and remaining 699 (12%) transcripts showed unique appearance to sporozoites invaded salivary glands (Fig. 5b). A heat map data analysis further suggested that salivary sporozoites may keep a strong hold on the salivary metabolic machinery, possibly to favor its own survival.

**Figure 5:**
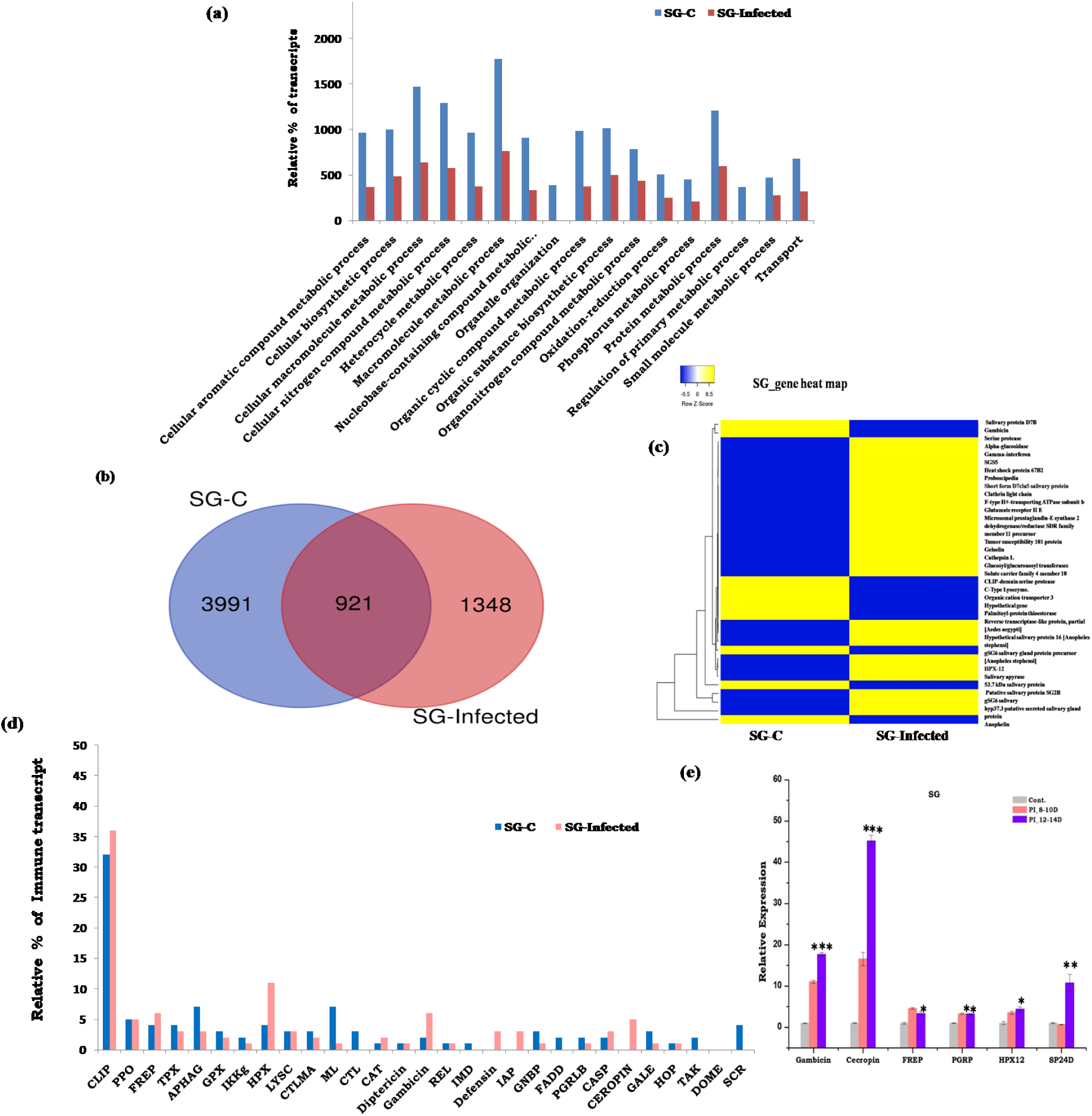
Detailed molecular map and comparative analysis of Blood fed (SG-C) and *P. vivax* infected (SG-Infected) salivary gland transcriptomes: **(a)** Gene ontology (GO-biological process/Level 4) based molecular cataloguing and comparison of salivary gland transcriptome in response to *P. vivax* infection; **(b)** Venn diagram analysis showing number of common and uncommon salivary transcripts altered in response to infection; **(c)** FPKM based heat map analysis expression of common salivary genes; **(d)** Molecular cataloguing of the immune genes enriched in the salivary glands of *Plasmodium* infected mosquitoes; **(e)** Transcriptional profiling of selected salivary gland genes (Gambicin p<0.0003; Cecropin p<0.0004; FREP: Fibrinogen related Proteins p<0.02; PGRP: Peptidoglycan Recognition Proteins p<0.002; HPX12: Heme peroxidase 12 p<0.001; SP24D: Serine Protease 24D p<0.09) during *P. vivax* infection(S2_primer list)

Next to test how salivary immune system influence sporozoites development and survival, we catalogued and compared relative percentage of immune transcripts (ST2). We noticed an exclusive appearance of four classes of immune family proteins such as FADD, Gambicin, GNBP and SCRC in the invaded salivary glands (Fig. 5d). While expression of TAK1, CASP, HOP and STAT immune family proteins remains restricted to naïve mosquito salivary glands (Fig. 5d). Transcriptional profiling of selected immune transcripts such as Gambicin, Cecropin, and SP-24 D showed a high induction in the salivary glands (Fig. 5e). Surprisingly, several other salivary secretory proteins such as Anopheline, D7 Family, Ion transporter family proteins, and 53.7kDa also showed a significant up-regulation suggesting their anti-plasmodium role against *P. vivax* infection.

### Deep sequencing identifies tissue specific *P. vivax* transcripts

The above analysis allowed us to hypothesize that when *Plasmodium sp.* switches from one stage to another, it may follow a unique strategy to regulate tissue-specific metabolic and immuno-physiological responses. However, clarifying the molecular nature of crosstalk between mosquito tissues and *Plasmodium* parasite remains a challenge. Surprising finding of a large pool of 4,449 transcripts of *P. vivax* origin, distinctly expressing in their respective tissues (Table-2), encouraged us to establish a genetic relationship of mosquito-parasite interaction. In fact, we noticed that 73.8% of *P. vivax* transcripts are associated with midgut oocysts (Early and late), while 74 transcripts are originating from free circulatory sporozoites in the hemolymph and 1,133 transcripts from salivary invaded Sporozoites(ST3).

**Table 2:**
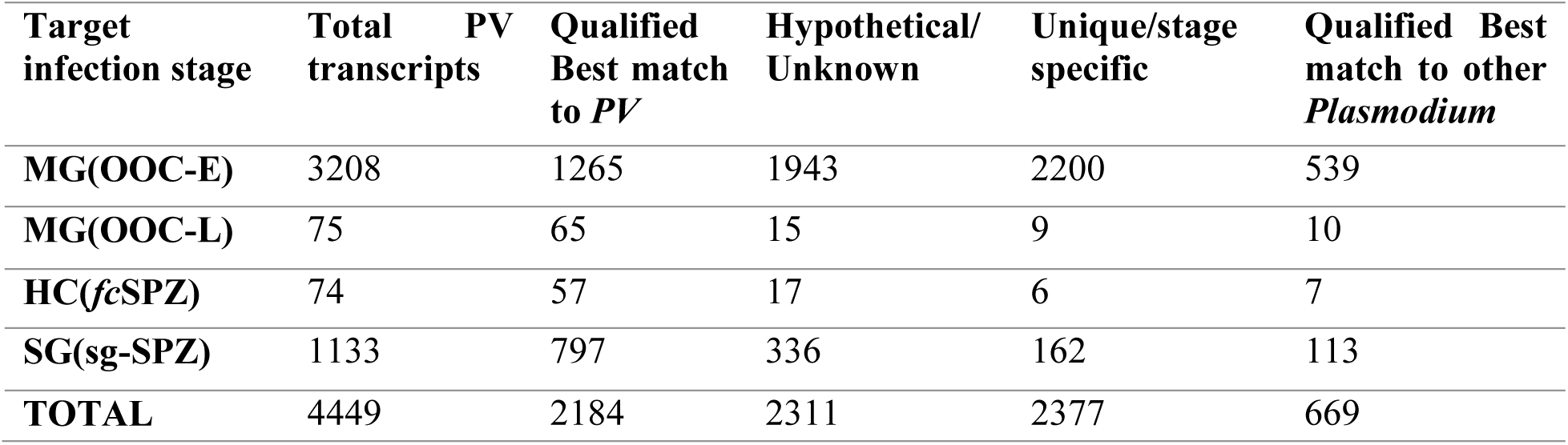
Detailed information of *P. vivax* transcripts of different life stages retrieved from mosquito transcriptomes.

### Molecular changes of *P. vivax* facilitate its survival and transmission

To establish a functional correlation, we annotated, catalogued and compared relative abundances of the *P. vivax* specific transcripts. Initially, putative transcripts encoding *Plasmodium* homolog proteins were filtered out from the pre analyzed RNAseq dataset. Each developmental stage of *P. vivax* revealed a large pool of hypothetical proteins especially in early oocysts and salivary sporozoites (Table-2), suggesting a complex biology is yet to unravel. Further, a GO annotation revealed that each developmental stage carries an enriched transcript abundance of the genes linked to cellular as well as metabolic processes (Fig.6a). Surprisingly, free circulatory sporozoites not only showed a selective enrichment for the family proteins encoding ‘Response to Stimulus’ and ‘Localization’, but also unraveled an exclusive appearance of signaling and detoxification linked genes in the hemocytes.

Since, rupturing of late oocysts release millions of *fc*SPZ which are highly vulnerable to hemocyte cellular immune response, we hypothesize that *Plasmodium* must have unique strategy to defend its survival. An in-depth functional annotation analysis (BP-Level 4), clearly demonstrated that despite of having low number of transcripts, late oocysts and *fc*SPZ encoded more diverse nature of proteins than early oocysts and salivary sporozoites (Fig. 6b). A Venn diagram analysis further unravel that the molecular nature of *Plasmodium* encoded proteins significantly altered when it switches from one stage to another, especially late oocysts (LO) to *fc*SPZ.

**Figure 6:**
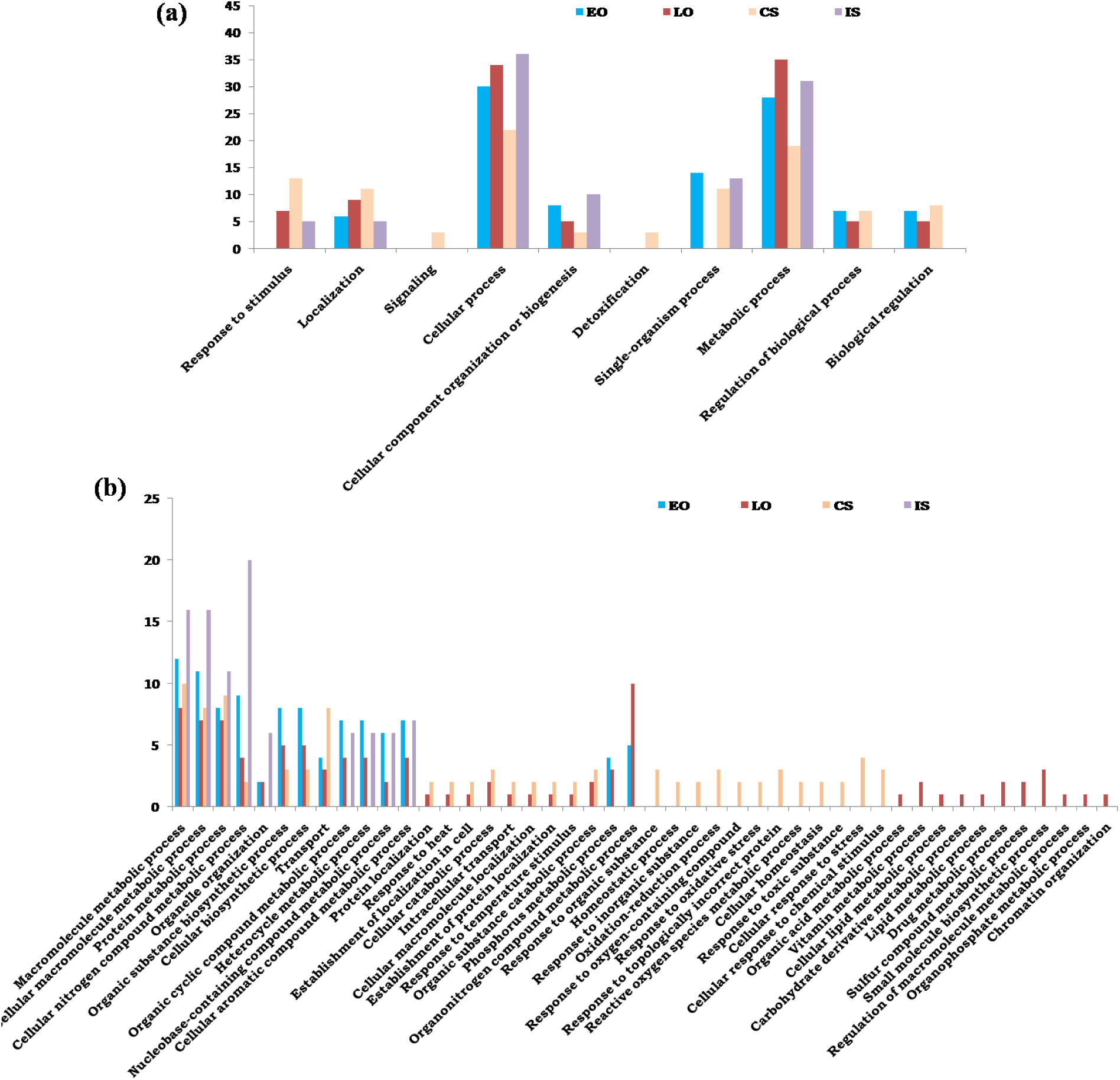
Molecular architecture of *P. vivax* transcripts during tissue specific developmental stages within the mosquito *An. stephensi*: **(a)** BLAST2GO (GO-biological process Level 2) based annotation and functional classification of identified *P. vivax* transcripts; **(b)** Detailed molecular and functional cataloguing (GO-biological process Level 4) of *P. vivax* genes reveal that, despite of having limited number of proteins, Late oocyst (LO) and free Circulatory sporozoite (CS) proteins are more diverse than Early oocyst (EO) and salivary Invaded sporozoite (IS) proteins.

Remarkably, a cross tissue comparison showed a restricted expression of large number of proteins in each developmental stage of *P. vivax* (Fig. 7a-c). Interestingly, a total of 2,200 transcripts showed restricted expression to early oocysts (EO), while 9 transcripts expression exclusively limited to late oocysts (LO) stage. Out of total 74 transcripts at least 6 showed restricted expression to *fc*SPZ stage and 162 transcripts expression exclusively remains to salivary invaded sporozoites (Fig. 7d). Though the exact mechanism of this developmental transformation is yet unknown. But a FPKM based relative expression analysis and transcriptional profiling of selected common transcripts further suggested that *P. vivax* may evolve with a unique ability of genetic makeup changes, possibly to misguide the immuno-physiological responses of mosquito tissues (Fig7e-f).

**Figure7:**
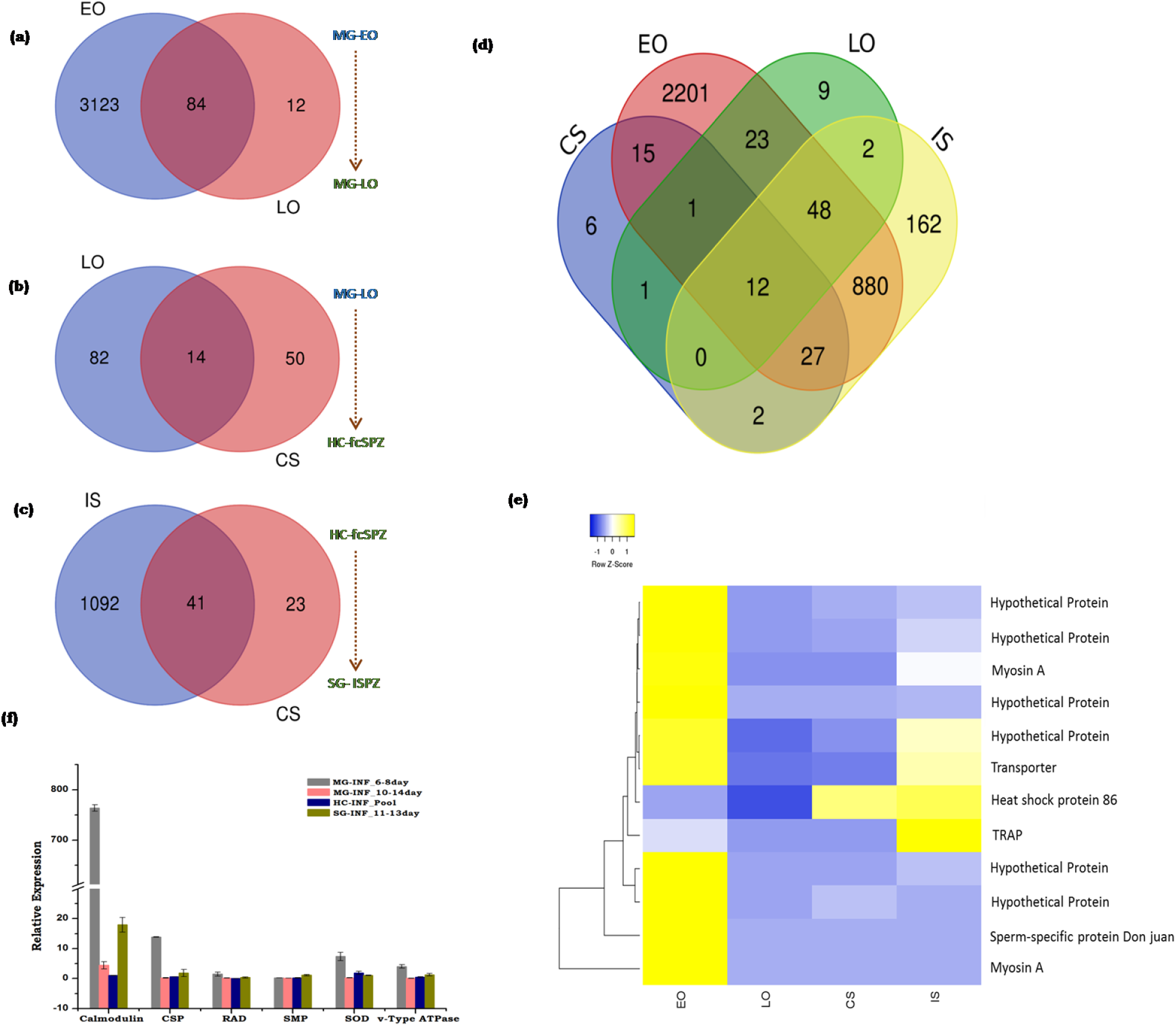
Stage specific comparative analysis of *P. vivax* transcripts: Venn diagrams showing stage specific *RNA-Seq* comparison between **(a)** Midgut Early oocyst (MG-EO) and Late oocyst (MG-LO); **(b)** Midgut Late oocyst (MG-LO) and free circulatory Sporozoite (HC-*fc*SPZ) of hemocytes and **(c)** HC-*fc*SPZ and salivary gland sporozoite (*sg*SPZ); (d) Venn diagram showing number of common and uncommon *P. vivax* transcripts expressing during distinct developmental stages i.e. Early oocyst (EO), Late oocyst (LO), free circulatory Sporozoite (*CS*) and salivary invaded Sporozoite (*IS*); (e) FPKM based comparative heat map of selected genes like myosin A, Transporter, TRAP (Throbospondin related adhesion protein) and some hypothetical proteins; **(f)** Tissue specific relative expression analysis of *P. vivax* genes (Calmodulin; CSP: Circumsporozoite protein; RAD protein; SMP: Sporozoite microneme protein; SOD: superoxide dismutase; vATPase).

Finally, we targeted to catalogue well annotated key genes which may have potential role in the parasite development and transmission. To do this we performed a comprehensive literature search and shortlisted transcripts with high FPKM, but restricted expression to a particular tissue (Fig. 8). Though, an ongoing independent detail comparison of *P. vivax* transcript database with other parasites such as *P. falciparum* and *P. berghei*, yet to unravel genetic differences (unpublished), current findings provide a valuable resource of *P. vivax* transcripts, especially free circulatory sporozoites directly interacting with mosquito hemocytes for future functional characterization.

**Figure 8:**
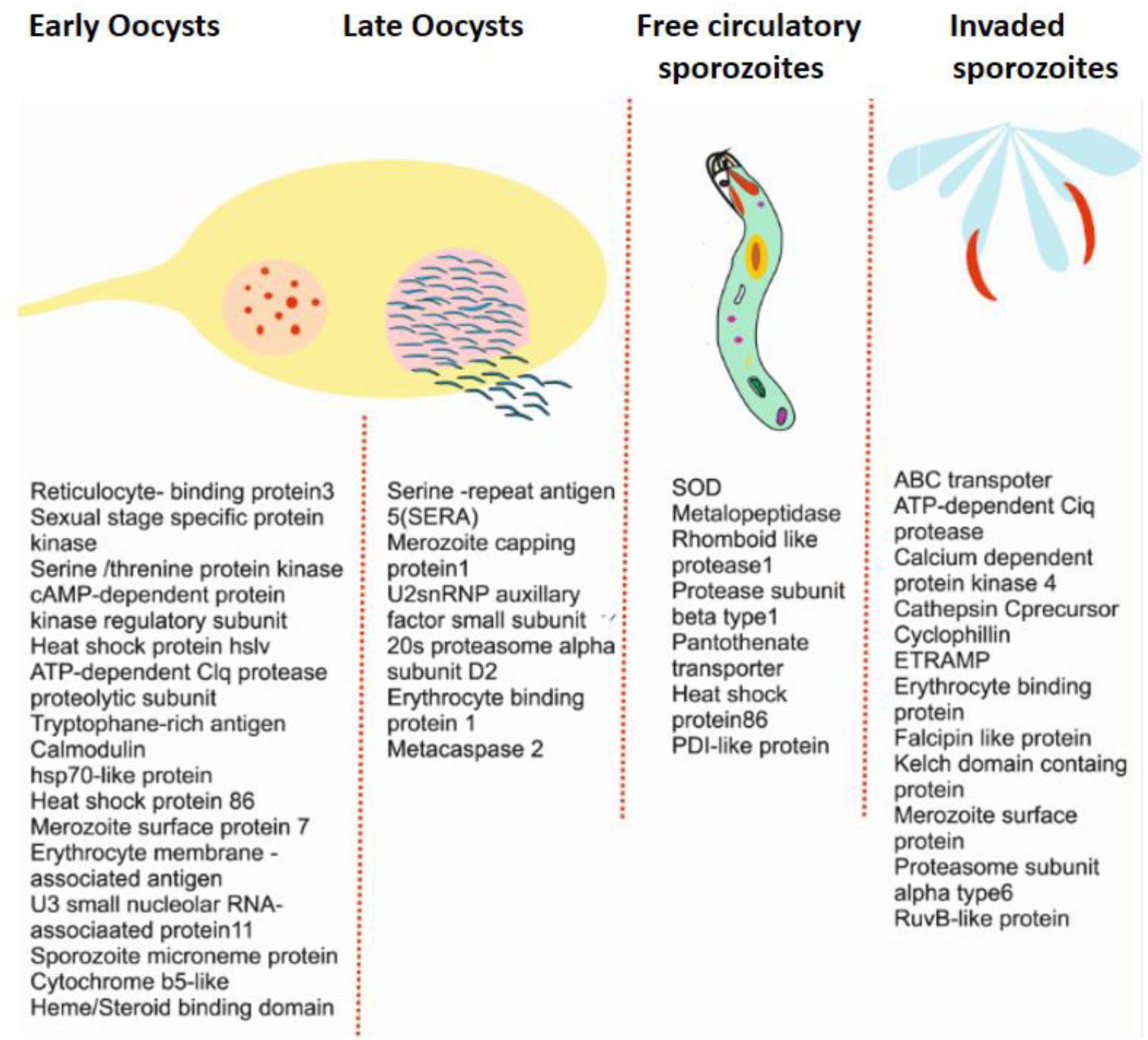
Representative catalogue of *RNAseq* identified *P. vivax* specific transcripts dominantly expressing in distinct mosquito tissues

## Discussion

Evolution and adaptation to blood feeding is an essential requirement for mosquitoes’ reproductive success (Attardo, Hansen, & Raikhel, 2005; Hansen, Attardo, Rodriguez, & Drake, 2014; Kokoza et al., 2001). *Plasmodium* parasites take the benefit of mosquitoes’ blood feeding behavior for the completion of sexual cycle and disease transmission (Thomas, 2012; Lacroix, Mukabana, Gouagna, & Koella, 2005; Schwartz & Koella, 2001). Thus, to win the developmental race, we opined that *Plasmodium* must have ability to manipulate immuno-physiological responses of directly interacting tissues such as midgut, hemocytes and salivary glands of mosquitoes (Cator et al., 2012). It has been well established that during its development inside the susceptible mosquito hosts, the population dynamics of *Plasmodium* significantly altered (Vézilier, Nicot, Gandon, & Rivero, 2012). But the nature of tissue specific molecular interactions, especially hemocyte-*Plasmodium* is not fully understood (Molina-Cruz et al., 2012). To decode this molecular complexity of mosquito-parasite interaction, we carried out a system-wide RNAseq analysis of each three tissue together. An initial profiling of AMPs/non-AMPs showed transient influence in tested tissue, suggesting a general role in the maintenance of the physiological homoeostasis (Fig. S4;).

Surprisingly, when compared to uninfected counterpart, each tissue shows a unique relationship of metabolic alteration in response to *P. vivax* infection. Our data demonstrated that post gut invasion, mosquito has the ability to regulate and recover from the acute harms caused by fast developing early oocysts. Many naturally selected refractory *Anopheline* species are able to cease the development of parasite through melanization (Molina-Cruz et al., 2012; Simões, Mlambo, Tripathi, Dong, & Dimopoulos, 2017). It is plausible to predict that even in the susceptible strain the controlled regulation of gut-parasite interplay is key for the survival of both mosquito and parasite (Dong, Manfredini, & Dimopoulos, 2009). Studies also suggest that a nutritional deprivation may have direct impact on the gut oocysts development and mosquitoes reproductive outcome, though mechanism is yet unknown (Liu, Dong, Huang, Rasgon, & Agre, 2013). Our observation of an early induction of transcripts regulating gut specific nutritional homeostasis such as Folliculin, Trehalase, Sterol carrier suggested that maturing oocysts are able to quench host nutritional resources (Baba et al., 2006; Schekman, 2013; Shukla, Thorat, Nath, & Gaikwad, 2015). However, delayed elevation of gut immune transcripts may counterbalance the negative impact of rapidly exiting sporozoites into the hemolymph. Thus, disrupting this relationship may favor the development of new tools to numb the parasites either by delimiting the nutritional demand, and/or enhancing the gut immunity (Shea-Donohue, Qin, & Smith, 2017).

Once left the gut epithelium, millions of *fc*SPZ directly encounter and cleared by hemocytes, but the molecular mechanism has not been fully established (Jullán F Hillyer, Schmidt, & Christensen, 2003; Jaramillo-Gutierrez et al., 2009). Our data suggests a major shift in transcripts abundance, especially encoding structural and catabolic activity associated proteins, may favor homeostasis maintenance of infected hemocytes. Transcriptional profiling further demonstrated that a hyper immunity is essential for majority of sporozoite population clearance. Several previous studies suggested that both REL and NO are key to regulate tissue specific immune regulation, and very recently we demonstrated that though both REL and NO participate inter-organ immune communication, but each tissue specifically maintains the inter-organ flow of signals (Clayton et al. 2014; Simões et al., 2017; Das De et al., 2018).

An indirect anti-*Plasmodium* response via hemocyte, REL immune signaling have also been suggested in the *An. gambiae, P. berghei* and *P. falciparum* model, but how hemocytes directly fights with *fc*SPZ is unknown (Julián F. Hillyer & Estévez-Lao, 2010). Our observation of increased REL than NOS against early *P. vivax* oocysts, further supported an idea that a pre-immune activation of hemocyte may exist (De et al., 2018; Kwon & Smith, 2018). But, surprisingly, still it remains unclear that how a fraction of *fc*SPZ succeed to avoid hyper immune response of hemocytes and invade salivary glands.

Our data on salivary-parasite interaction indicated *Plasmodium* sporozoites not only impair the metabolic machinery, but also enriches nucleic acid binding transcriptional activities. For a successful blood meal acquisition, salivary glands releases a cocktail containing nucleic acid binding factors such as Apyrase (King, Vernicks, & Hillyer, 2011), D7 family proteins (Calvo, Mans, Andersen, & Ribeiro, 2006) and nucleotide transferase (Dhar & Kumar, 2003; Vogt et al., 2018). Thus it is plausible to hypothesize that *Plasmodium* infection may enhance host seeking behavioral activities by stimulating the expression of nucleic acid binding factors (Chen, Mathur, & James, 2008). Additionally, an increased expression of salivary immune transcripts indicated that an active local immune response is essential to restrict salivary invaded sporozoite (IS) population.

As a proof of concept, our data suggested that even after mounting an effective tissue specific immuno-physiological response, mosquitoes fail to disrupt *Plasmodium* sporogonic cycle. Hence, we hypothesize that *Plasmodium* parasites are clever enough to dodge the mosquito’s immune system by wise manipulation of its own molecular architecture. Retrieval of large pool of *P. vivax* transcripts originating from distinct developmental stages provided us an opportunity to anticipate the molecular dynamics facilitating its survival. An initial observation of more than 50% transcripts encoding hypothetical proteins indicated that a deep understanding of *P. vivax* sporogonic cycle in still obscure. We also observed a significant difference in the molecular repertoire of stage specific *P. vivax* genes further strengthen our hypothesis that *Plasmodium* parasite has unique ability to misguide the track and trap system of mosquitoes.

## Conclusion

For its successful survival and transmission, every stage of development inside the mosquito host, *Plasmodium* negotiates multiple tissues. Independent studies, targeting individual tissues have been valuable, but conceptually we have unresolved major questions that how mosquito’s tissue specific actions manage the challenge of *Plasmodium* infection and/or how *Plasmodium* manages to avoid the tissue specific responses. For the first time we demonstrate and establish that *P. vivax* follows a smart strategy of genetic makeup change to misguide and evade, even a highly sophisticated immune barrier of each tissue. We hypothesize that an unharmed tissue specific molecular wave of negotiations and actions by genetic changes benefits *P. vivax* successful development and transmission in the mosquito host (Fig 9). Further establishing a functional correlation may lead to identification of mosquito as well as *P. vivax* specific crucial genetic factors for target selection and designing new molecular strategy for malaria control.

**Figure9:**
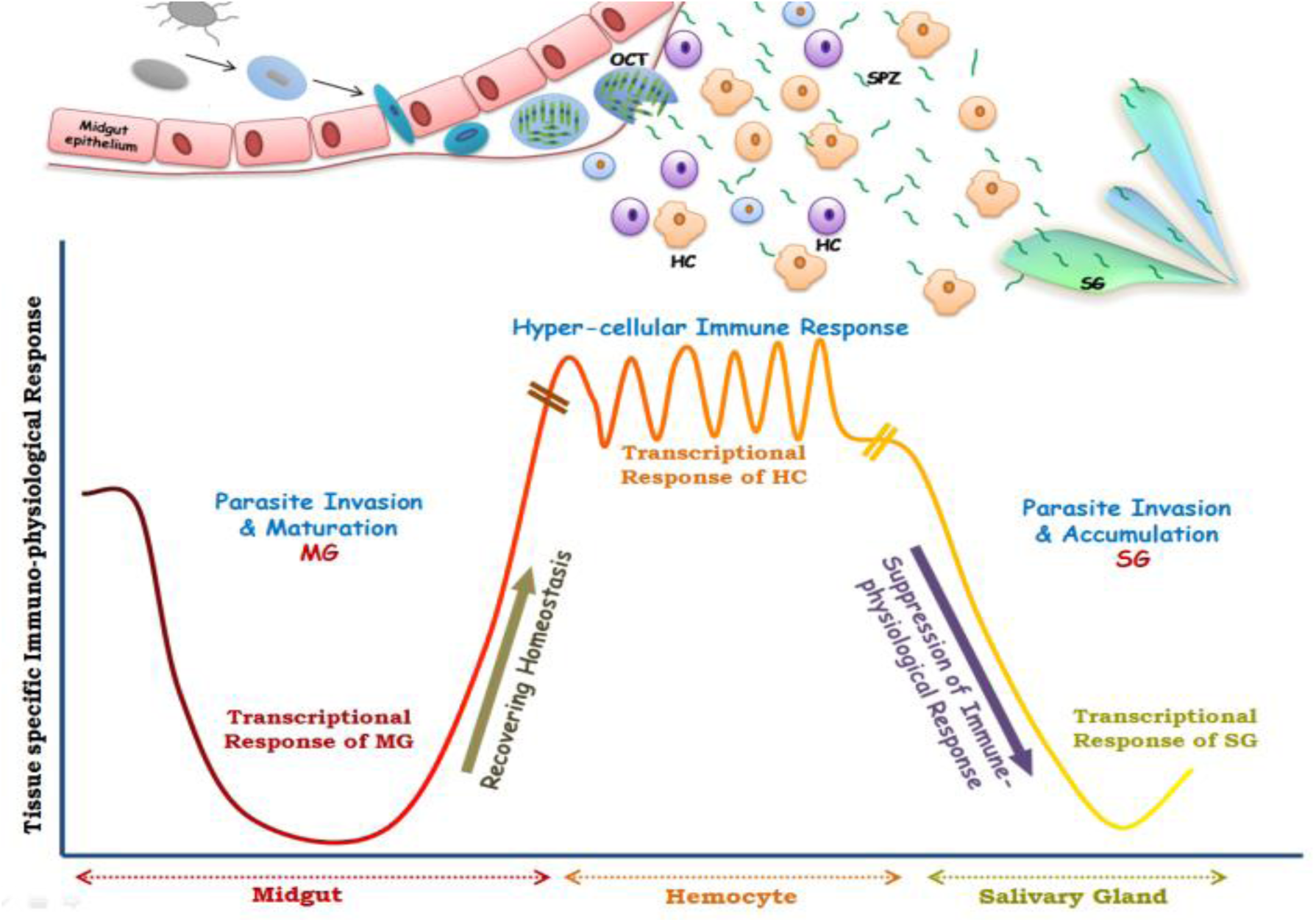
Proposed future working hypothesis to understand tissue specific molecular interactions during complete development of *P. vivax* in the mosquito host: As the *Plasmodium* parasites enter the midgut (MG) through infected blood meal, it undergoes fertilization and form zygotes, which further transform into ookinetes. These ookinetes then invade the midgut epithelium and reside within the basal lamina, where they form oocysts (OCT) and get matured. During this period midgut metabolic machinery remains suppressed to fulfill parasites’ nutritional demand and as the parasite leaves the basal lamina in the form of free circulatory sporozoites (SPZ), MG starts regaining its homeostasis. After sporozoites get perfuse in mosquito hemolymph they have to interact directly with hemocytes (HC) which shows hyper cellular immune response to delimit parasite population. Survived sporozoites proceed to invade the salivary glands and accumulate there for further transmission to next host, and during this period salivary gland transcriptional response remain suppressed. Ability of *Plasmodium vivax* to alter developmental stage specific genetic changes may help to evade tissue specific mosquito response, favoring successful survival and transmission.

## Supporting information

Immune catalogue

PV_transcripts_catalogue

RNAseq_Annotation

## DATA DEPOSITION

The sequences of the individual tissue sample were submitted to the NCBI having accession numbers SRR8580009 (MG-Inf 8-10D); SRR8580010 (MG-C); SRR8580011 (MG-Inf 3-6D); SRR8501353 (HC-C); SRR8501352 (HC-PI); SRR8476334 (SG-PI); SRR8476333 (SG-C).

## ACKNOWLEDGEMENT

We would like to thank all the technical staff members of central insectary facility for mosquito rearing and Kunverjeet singh for lab assistance. Authors thanks to NIMR-clinical facility support. Finally, we thank Xceleris Genomics, Ahmedabad for NGS sequencing.

## Authors’ contribution

ST, SK, CC, KCP, VP, RKD conceived scientific hypothesis and designed the experiments, ST, SK, CC performed tissue i.e. midgut, salivary gland and hemocyte specific experiments, DS, PS, TT, JR, TDD contributed to design and performing the experiments, data acquisition, writing and editing; ST, SK, CC, KCP, VP, RKD data analysis and interpretation, data presentation, contributed reagents/ materials/Analysis tools, wrote, reviewed, edited, and finalized MS. All authors read and approved the final manuscript

## References

Aly, A. S. I., Vaughan, A. M., & Kappe, S. H. I. (2009). Malaria Parasite Development in the Mosquito and Infection of the Mammalian Host. Annual Review of Microbiology. https://doi.org/10.1146/annurev.micro.091208.073403

Attardo, G. M., Hansen, I. A., & Raikhel, A. S. (2005). Nutritional regulation of vitellogenesis in mosquitoes: Implications for anautogeny. In Insect Biochemistry and Molecular Biology. https://doi.org/10.1016/j.ibmb.2005.02.013

Baba, M., Hong, S.-B., Sharma, N., Warren, M. B., Nickerson, M. L., Iwamatsu, A., … Zbar, B. (2006). Folliculin encoded by the BHD gene interacts with a binding protein, FNIP1, and AMPK, and is involved in AMPK and mTOR signaling. Proceedings of the National Academy of Sciences, 103(42), 15552–15557. https://doi.org/10.1073/pnas.0603781103

Baton, L. A., & Ranford-Cartwright, L. C. (2004). *Plasmodium falciparum* ookinete invasion of the midgut epithelium of Anopheles stephensi is consistent with the Time Bomb model. Parasitology. https://doi.org/10.1017/s0031182004005979

Belachew, E. B. (2018). Immune Response and Evasion Mechanisms of Plasmodium falciparum Parasites. Journal of Immunology Research. https://doi.org/10.1155/2018/6529681

Bennink, S., Kiesow, M. J., & Pradel, G. (2016). The development of malaria parasites in the mosquito midgut. Cellular Microbiology. https://doi.org/10.1111/cmi.12604

Calvo, E., Mans, B. J., Andersen, J. F., & Ribeiro, J. M. C. (2006). Function and evolution of a mosquito salivary protein family. Journal of Biological Chemistry. https://doi.org/10.1074/jbc.M510359200

Castillo, J. C., Barletta Ferreira, A. B., Trisnadi, N., & Barillas-Mury, C. (2017). Activation of mosquito complement antiplasmodial response requires cellular immunity. Science Immunology. https://doi.org/10.1126/sciimmunol.aal1505

Cator, L. J., Lynch, P. A., Read, A. F., & Thomas, M. B. (2012). Do malaria parasites manipulate mosquitoes? Trends in Parasitology. https://doi.org/10.1016/j.pt.2012.08.004

Chen, X. G., Mathur, G., & James, A. A. (2008). Chapter 2 Gene Expression Studies in Mosquitoes. Advances in Genetics. https://doi.org/10.1016/S0065-2660(08)00802-X

Cirimotich, C. M., Dong, Y., Garver, L. S., Sim, S., & Dimopoulos, G. (2010). Mosquito immune defenses against Plasmodium infection. Developmental and Comparative Immunology. https://doi.org/10.1016/j.dci.2009.12.005

Clayton, A. M., Dong, Y., & Dimopoulos, G. (2014). The anopheles innate immune system in the defense against malaria infection. Journal of Innate Immunity. https://doi.org/10.1159/000353602

Conesa, A., Götz, S., García-gómez, J. M., Terol, J., & Talón, M. (2005). Sequence analysis Blast2GO: a universal tool for annotation, visualization and analysis in functional genomics research, (October), 2–5. https://doi.org/10.1093/bioinformatics/bti610

De, T. Das, Sharma, P., Thomas, T., Singla, D., Tevatiya, S., Kumari, S., … Dixit, R. (2018). Interorgan molecular communication strategies of “Local” and “Systemic” innate immune responses in mosquito Anopheles stephensi. Frontiers in Immunology. https://doi.org/10.3389/fimmu.2018.00148

Dhar, R., & Kumar, N. (2003). Role of mosquito salivary glands. Current Science.

Dixit, R., Rawat, M., Kumar, S., Pandey, K. C., Adak, T., & Sharma, A. (2011). Salivary gland transcriptome analysis in response to sugar feeding in malaria vector anopheles stephensi. Journal of Insect Physiology. https://doi.org/10.1016/j.jinsphys.2011.07.007

Dong, Y., Manfredini, F., & Dimopoulos, G. (2009). Implication of the mosquito midgut microbiota in the defense against malaria parasites. PLoS Pathogens. https://doi.org/10.1371/journal.ppat.1000423

Garver, L. S., de Almeida Oliveira, G., & Barillas-Mury, C. (2013). The JNK Pathway Is a Key Mediator of Anopheles gambiae Antiplasmodial Immunity. PLoS Pathogens. https://doi.org/10.1371/journal.ppat.1003622

Gouagna, L. C., Mulder, B., Noubissi, E., Tchuinkam, T., Verhave, J. P., & Boudin, C. (1998). The early sporogonic cycle of Plasmodium falciparum in laboratory-infected Anopheles gambiae: An estimation of parasite efficacy. Tropical Medicine and International Health. https://doi.org/10.1046/j.1365-3156.1998.00156.x

Han, Y. S., Thompson, J., Kafatos, F. C., & Barillas-Mury, C. (2000). MC8 - Molecular interactions between Anopheles stephensi midgut cells and Plasmodium berghei: The time bomb theory of ookinete invasion. Memorias Do Instituto Oswaldo Cruz. https://doi.org/10.1093/emboj/19.22.6030

Hansen, I. A., Attardo, G. M., Rodriguez, S. D., & Drake, L. L. (2014). Four-way regulation of mosquito yolk protein precursor genes by juvenile hormone-, ecdysone-, nutrient-, and insulin-like peptide signaling pathways. Frontiers in Physiology. https://doi.org/10.3389/fphys.2014.00103

Hillyer, Julián F., & Estévez-Lao, T. Y. (2010). Nitric oxide is an essential component of the hemocyte-mediated mosquito immune response against bacteria. Developmental and Comparative Immunology. https://doi.org/10.1016/j.dci.2009.08.014

Hillyer, Julián F., Schmidt, S. L., & Christensen, B. M. (2003). Hemocyte-mediated phagocytosis and melanization in the mosquito Armigeres subalbatus following immune challenge by bacteria. Cell and Tissue Research. https://doi.org/10.1007/s00441-003-0744-y

Hillyer, Jullán F, Schmidt, S. L., & Christensen, B. M. (2003). Rapid phagocytosis and melanization of bacteria and Plasmodium sporozoites by hemocytes of the mosquito Aedes aegypti. The Journal of Parasitology. https://doi.org/10.1645/0022-3395(2003)089[0062:RPAMOB]2.0.CO;2

Jaramillo-Gutierrez, G., Rodrigues, J., Ndikuyeze, G., Povelones, M., Molina-Cruz, A., & Barillas-Mury, C. (2009). Mosquito immune responses and compatibility between Plasmodium parasites and anopheline mosquitoes. BMC Microbiology. https://doi.org/10.1186/1471-2180-9-154

Kariu, T., Yuda, M., Yano, K., & Chinzei, Y. (2002). MAEBL Is Essential for Malarial Sporozoite Infection of the Mosquito Salivary Gland. The Journal of Experimental Medicine. https://doi.org/10.1084/jem.20011876

King, J. G., Vernicks, K. D., & Hillyer, J. F. (2011). Members of the salivary gland surface protein (SGS) family are major immunogenic components of mosquito saliva. Journal of Biological Chemistry. https://doi.org/10.1074/jbc.M111.280552

Kokoza, V. A., Martin, D., Mienaltowski, M. J., Ahmed, A., Morton, C. M., & Raikhel, A. S. (2001). Transcriptional regulation of the mosquito vitellogenin gene via a blood meal-triggered cascade. Gene. https://doi.org/10.1016/S0378-1119(01)00602-3

Kuehn, A., & Pradel, G. (2010). The Coming-Out of Malaria Gametocytes. Journal of Biomedicine and Biotechnology. https://doi.org/10.1155/2010/976827

Kumar, S., Gupta, L., Yeon, S. H., & Barillas-Mury, C. (2004). Inducible peroxidases mediate nitration of Anopheles midgut cells undergoing apoptosis in response to Plasmodium invasion. Journal of Biological Chemistry. https://doi.org/10.1074/jbc.M409905200

Kwon, H., & Smith, R. C. (2018). Chemical depletion of phagocytic immune cells reveals dual roles of mosquito hemocytes in Anopheles gambiae anti-Plasmodium 2 immunity 3. https://doi.org/10.1101/422543

Lacroix, R., Mukabana, W. R., Gouagna, L. C., & Koella, J. C. (2005). Malaria infection increases attractiveness of humans to mosquitoes. PLoS Biology. https://doi.org/10.1371/journal.pbio.0030298

Lam, F., Lalansingh, C. M., Babaran, H. E., Wang, Z., Prokopec, S. D., Fox, N. S., & Boutros, P. C. (2016). VennDiagramWeb: A web application for the generation of highly customizable Venn and Euler diagrams. BMC Bioinformatics. https://doi.org/10.1186/s12859-016-1281-5

Liu, K., Dong, Y., Huang, Y., Rasgon, J. L., & Agre, P. (2013). Impact of trehalose transporter knockdown on Anopheles gambiae stress adaptation and susceptibility to Plasmodium falciparum infection. Proceedings of the National Academy of Sciences. https://doi.org/10.1073/pnas.1316709110

Molina-Cruz, A., DeJong, R. J., Ortega, C., Haile, A., Abban, E., Rodrigues, J., … Barillas-Mury, C. (2012). Some strains of Plasmodium falciparum, a human malaria parasite, evade the complement-like system of Anopheles gambiae mosquitoes. Proceedings of the National Academy of Sciences. https://doi.org/10.1073/pnas.1121183109

Mueller, A. K., Kohlhepp, F., Hammerschmidt, C., & Michel, K. (2010). Invasion of mosquito salivary glands by malaria parasites: Prerequisites and defense strategies. International Journal for Parasitology. https://doi.org/10.1016/j.ijpara.2010.05.005

Ramphul, U. N., Garver, L. S., Molina-Cruz, A., Canepa, G. E., & Barillas-Mury, C. (2015). Plasmodium falciparum evades mosquito immunity by disrupting JNK-mediated apoptosis of invaded midgut cells. Proceedings of the National Academy of Sciences. https://doi.org/10.1073/pnas.1423586112

Roth, A., Adapa, S. R., Zhang, M., Liao, X., Saxena, V., Goffe, R., … Adams, J. H. (2018). Unraveling the Plasmodium vivax sporozoite transcriptional journey from mosquito vector to human host. Scientific Reports. https://doi.org/10.1038/s41598-018-30713-1

Schekman, R. (2013). Discovery of the cellular and molecular basis of cholesterol control. Proceedings of the National Academy of Sciences, 110(37), 14833–14836. https://doi.org/10.1073/pnas.1312967110

Schwartz, A., & Koella, J. C. (2001). Trade-offs, conflicts of interest and manipulation in Plasmodium-mosquito interactions. Trends in Parasitology. https://doi.org/10.1016/S1471-4922(00)01945-0

Sharma, P., Sharma, S., Mishra, A. K., Thomas, T., Das De, T., Rohilla, S. L., … Dixit, R. (2015). Unraveling dual feeding associated molecular complexity of salivary glands in the mosquito Anopheles culicifacies. Biology Open. https://doi.org/10.1242/bio.012294

Shea-Donohue, T., Qin, B., & Smith, A. (2017). Parasites, nutrition, immune responses and biology of metabolic tissues. Parasite Immunology. https://doi.org/10.1111/pim.12422

Shiao, S. H., Whitten, M. M. A., Zachary, D., Hoffmann, J. A., & Levashina, E. A. (2006). Fz2 and Cdc42 mediate melanization and actin polymerization but are dispensable for Plasmodium killing in the mosquito midgut. PLoS Pathogens. https://doi.org/10.1371/journal.ppat.0020133

Shukla, E., Thorat, L. J., Nath, B. B., & Gaikwad, S. M. (2015). Insect trehalase: Physiological significance and potential applications. Glycobiology. https://doi.org/10.1093/glycob/cwu125

Simões, M. L., Mlambo, G., Tripathi, A., Dong, Y., & Dimopoulos, G. (2017). Immune Regulation of Plasmodium Is Anopheles Species Specific and Infection Intensity Dependent. MBio. https://doi.org/10.1128/mbio.01631-17

Smith, R. C., King, J. G., Tao, D., Zeleznik, O. A., Brando, C., Thallinger, G. G., & Dinglasan, R. R. (2016). Molecular Profiling of Phagocytic Immune Cells in *Anopheles gambiae* Reveals Integral Roles for Hemocytes in Mosquito Innate Immunity. Molecular & Cellular Proteomics. https://doi.org/10.1074/mcp.M116.060723

Thomas, T., De, D.T., Sharma, P., Sharma, S., Lata, S., Saraswat, P., Pandey, K.C., Dixit, R. (2016). Hemocytome: deep sequencing analysis of mosquito blood cells in Indian malarial vector Anopheles stephensi. Gene. https://doi.org/10.1016/j.gene.2016.02.031

Talman, A. M., Domarle, O., McKenzie, F. E., Ariey, F., & Robert, V. (2004). Gametocytogenesis: The puberty of Plasmodium falciparum. Malaria Journal. https://doi.org/10.1186/1475-2875-3-24

Thomas, T., Das De, T., Sharma, P., Lata, S., Saraswat, P., Pandey, K. C., & Dixit, R. (2016). Hemocytome: Deep sequencing analysis of mosquito blood cells in Indian malarial vector Anopheles stephensi. Gene. https://doi.org/10.1016/j.gene.2016.02.031

Vézilier, J., Nicot, A., Gandon, S., & Rivero, A. (2012). Plasmodium infection decreases fecundity and increases survival of mosquitoes. Proceedings of the Royal Society B: Biological Sciences. https://doi.org/10.1098/rspb.2012.1394

Vogt, M. B., Lahon, A., Arya, R. P., Kneubehl, A. R., Spencer Clinton, J. L., Paust, S., & Rico-Hesse, R. (2018). Mosquito saliva alone has profound effects on the human immune system. PLoS Neglected Tropical Diseases. https://doi.org/10.1371/journal.pntd.0006439

