## Supplemental Data File for "Genetic changes of *P. vivax* tempers host tissue-specific responses in *Anopheles stephensi*"

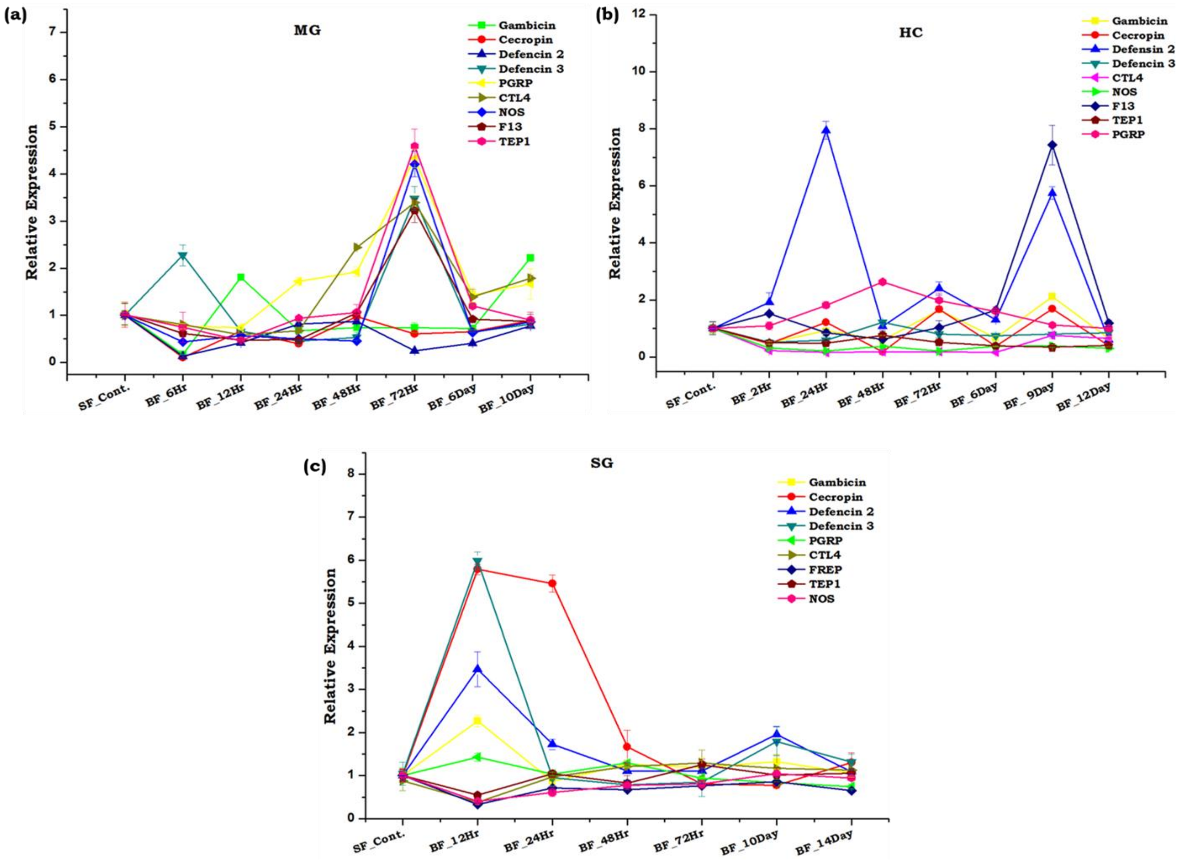


**Figure S1**: **Tissue specific differential modulation of immune gene following blood meal uptake:**Relative gene expression analysis of Anti microbial peptides (AMPs: Gambicin, Defencin 2,Defencin 3, Cecropin)and Non-AMPs (F13:Fibrinogen related protein 13; PGRP:Peptidoglycan Recognition Proteins; CTL4:C-type lectin 4; TEP1:Thioester-containing protein 1; NOS: Nitric oxide synthase)in response to blood feeding in (a)Midgut (MG) (b) Hemocytes (HC) and (c) Salivary Gland (SG)

### Table S2: List of primers used for the study

| | Gene Name | Primer Sequence |
|---|---|---|
| 1 | Cecropin | Fw: AACCAACCGAACCGTATCAA |
| | | Rev: ACCGAAACCAGAACAAATCG |
| 2 | Defensin | Fw: GCGACCGTGTGTAGTTCAGA |
| | | Rev: CACTCATCCTGGTCGCTACA |
| 3 | Gambicin | Fw: ACTGTGGCTACGGGTACGTC |

| | | |
|---|---|---|
| | | Rev: GCTTGTTCTTCCGGTGTGAT |
| 4 | Heme peroxidase 12 | Fw: GAACAGTGCCACCGATACCT |
| | | Rev: CCGAGATAATAGGGCAACCA |
| 5 | Serine protease 24 | Fw: ACCGTGCTACCGTAATCCAG |
| | | Rev: GGCACATAGTCGGACACCTT |
| 6 | vacuolar ATP synthetase | Fw: CATAGTAGGCCTGAGTTCAT |
| | | Rev: GAGATGTAAATGCCGATAAT |
| 7 | RAD | Fw: GTATTTACAACGCCGTTACT |
| | | Rev: CTCGAAGAAATGTTTGTAGG |
| 8 | Sterol carrier | Fw: TGAAGGTGGAGAAAGGTACGA |
| | | Rev: TAGATGACTTCGGCCAGCTT |
| 9 | Folliculin | Fw: CTCGCTATTTGACAGAAAGC |
| | | Rev: GAAACATCCGCAGAGAGTAT |
| 10 | Trehalase | Fw: ACGTTTAACAAAACGAGAGC |
| | | Rev: GCCGGTATTGATGGAGTAT |
| 11 | Calmodulin | Fw: CAGACAAGTTAACGGAGGAG |
| | | Rev: CGTGTCCGTATCTTTCATTT |
| 12 | CSP | Fw: GGACAGGGACAAAATAATGA |
| | | Rev: ATATGCCAGCACACTTATCC |
| 13 | F13 | Fw: AGTTTGAGATAGGCAGCGA |
| | | Rev: CATCCACCGCATTCATGAA |
| 14 | Superoxide Dismutase (SOD) | Fw:GACACTTTGGATCAGGATGG |
| | | Rev:GGCAAAATTCCAGTTGACC |
| 15 | SporozoiteMicroneme Protein (SMP) | Fw:CCAGGCCACGGAATACTT |
| | | Rev:ATGACGCCTTTGCTGTTTT |
| 16 | ApolipophorinIII | Fw: CATCGTGAAGGATCTGAAAG |
| | | Rev: TGAAGACTTCGAAAGTCGAG |

| | | |
|---|---|---|
| 17 | LRIM 17 | Fw: ATCTGGATCTGAACGTGATG |
| | | Rev: AGATTCTGATTCTTGCGACA |
| 18 | Hexamerin | Fw: TATGCCGATGCAGTTCTACT |
| | | Rev: TGGTTCATTTTGATCTCGTC |
| 19 | Actin | Fw: TGCGTGACATCAAGGAGAAG |
| | | Rev: GATTCCATACCCAGGAACGA |
| 20 | Nitric oxide synthase | Fw: GACCAAACCGGTCATCCTGAT |
| | | Rev: GGAATCTTGCAGTCAACCATTTC |
